# Viability, task switching, and fall avoidance of the simplest dynamic walker

**DOI:** 10.1101/2022.01.16.476517

**Authors:** Navendu S. Patil, Jonathan B. Dingwell, Joseph P. Cusumano

**Affiliations:** Department of Kinesiology, Pennsylvania State University, University Park, PA 16802, USA; Department of Engineering Science & Mechanics, Pennsylvania State University, University Park, PA 16802, USA

## Abstract

Humans display great versatility when performing goal-directed tasks while walking. However, the extent to which such versatility helps with fall avoidance remains unclear. We recently demonstrated a functional connection between the motor regulation needed to achieve task goals (e.g. maintaining walking speed) and a simple walker’s ability to reject large disturbances. Here, for the same model, we identify the viability kernel—the state space region in which the walker can step forever via at least one sequence of push-off inputs per state. We further find that only a few basins of attraction of the speed-regulated walker’s steady-state gaits can fully cover the viability kernel. This highlights a potentially important role of task-level motor regulation in fall avoidance. Therefore, we posit an adaptive hierarchical control/regulation strategy that switches between different task-level regulators to avoid falls. Our hierarchical task switching controller only requires a target value of the regulated observable—a ‘task switch’—at each walking step, each chosen from a small, predetermined collection. Because humans have typically already learned to perform such tasks during nominal walking conditions, this suggests that the ‘information cost’ of biologically implementing such controllers for the nervous system, including cognitive demands in humans, could be relatively low.

## Introduction

When human infants learn to walk, they essentially learn, albeit after extensive practice, to be ‘viable’, i.e. to avoid falls^1^. Likewise, older adults frequently fall while walking, and the related injuries are a serious public health issue^2,3^. Quantifying individuals’ predisposition to fall is critical to minimizing fall incidence. However, the risk of a fall in humans likely depends on multiple biomechanical, neurological, cognitive, and environmental factors^4^. Stability while walking is not automatic, as even healthy humans need to actively cope with physiological motor noise and environmental disturbances. Here, we focus on the *global* dynamic stability of walking, i.e. a walker’s ability to reject *large* external disturbances (as might occur, e.g. while avoiding an unanticipated obstacle, or from a ‘shove’), which is central to avoiding falls.

The paradigm of ‘limit cycle walking’^5,6^ has shown that nominally periodic, disturbance-free walking motions (i.e. ‘limit cycles’) that are stable can be achieved without requiring continuous-time active control of walking trajectories between step transitions. Such nominal limb trajectories, across a variety of human movements, have also been predicted within the optimal control framework consistent with the ‘minimum intervention’ principle^7^. Many optimality-based approaches naturally seek solutions in the form of a single limit cycle, having a specific set of gait parameters such as step length and step time, often known *a priori*. However, such solutions are excessively restrictive as they overly constrain walking motion around a single trajectory. In contrast, humans have necessarily learned to walk at many such limit cycles, both stably and efficiently, to remain versatile. The notion of viability^8^ is well-suited to handle the non-uniqueness of solutions to a given walking task as it does not seek to optimize desired trajectories, but only to avoid falls. Moreover, at least in principle, humans could remain viable using control strategies that quickly switch between multiple limit cycles.

Typically, humans also walk to achieve one or more task goals, by maintaining one or more gait observables (i.e. empirically measurable variables like speed or direction). Moreover, any given walking task could be performed via multiple gait patterns, each specified by a set of gait parameters^9,10^. Such task-level redundancies also interact with biomechanical redundancies at the level of muscles and joints^11^. This non-uniqueness of solutions to a given motor task makes the problem of biological movement control mathematically ill-posed. For tasks that have multiple uncertain goals with similar costs, humans deliberately select intermediate movements to maximize task success^12^. In walking, however, fall avoidance decidedly supersedes achieving other task goals, which themselves often have different priorities^10^. This makes the choice of an optimal strategy far less obvious.

In this vein, our work is motivated by the following fundamental questions: How does the nervous system manage redundancies while achieving task goals in any given walking context? Furthermore, how might this functional organization help minimize fall incidence? In response, we posit that humans achieve stable and efficient walking gaits via a *hierarchical schema*, consisting of what we will here distinguish as *control* versus *regulation* of movement. Specifically, we use ‘control’ to refer to the processes required for a walker to remain *viable* while taking individual steps. Conversely, we use ‘regulation’ to refer to the processes needed to carry out specific goal-directed *tasks*. Evidently, the walker must remain viable at all times, including while carrying out specific walking tasks. Thus, control and regulation, while functionally distinct, are hierarchical by design.

Viability is a generalized form of stability of an actuated dynamical system such that the system can avoid failure forever by choosing appropriate sequence(s) of inputs within its actuator limits^8^. The set of viable states in the system’s state space, in which its viability is guaranteed, thus provides a measure of the system’s ability to avoid failure. Indeed, the bigger the viable set of states in a walker’s state space, the better is the walker’s ability to avoid falls, as it can, at least in principle, reject a larger range of external disturbances^13–15^. However, by itself, viability does not take into account goal-directedness in walking. Indeed, the walker could, at least in principle, step randomly (i.e. with no ‘intent’) forever within the viable region.

Task-level motor regulation allows the walker to achieve specific task goals by targeting relevant task-level observables from each walking step to the next: it rapidly corrects deviations in stepping observables that interfere with achieving a specific goal, while allowing task-irrelevant deviations to persist^9,16^. Thus, regulation, too, can permit walking at several limit cycles, as long as they do not violate the specific task requirements, especially while stability concerns are not paramount. However, by itself, task-level regulation does not aim to guarantee stability of the walker’s limit cycle, let alone maximize its global stability.

Our recent work^17^ highlights a possible answer as to why humans might prefer one equally workable task-level regulation strategy over another from the perspective of fall avoidance. We studied this question by integrating the simplest *mechanical template*^18^ of walking on level ground with *motor regulation templates*, i.e. empirically motivated models of how humans manipulate task-level observables on a step-to-step basis^10,16^. In experiments, humans walking on a treadmill tightly regulate speed at successive strides, while allowing absolute position to drift for many strides^9,16^. We simulated a push-off powered compass walker^19^ that additionally regulated step-to-step speed or absolute position on a treadmill. We characterized global stability of the walker’s limit cycles (i.e. steady-state periodic gaits) by the size and shape of their *basins of attraction* in the state space. Task-level regulation, despite not being designed to do so, makes walking more robust to external disturbances: it yields superior *local* (i.e. small) disturbance rejection and improved *global* stability, both by increasing the size of basins of attraction and by regularizing their geometric structure^17^. While both step-to-step speed and position regulation provide workable strategies for treadmill walking, speed regulation enlarges and regularizes the unregulated walker’s basin much more than position regulation. These simulation results are consistent with experiments^9,16^ and thus demonstrate a functional connection between task-level motor regulation and global stability. However, that prior work did not assess motor regulation strategies within the context of viability.

Here, we extend this recent work and study the same powered walker (Fig. 1) to identify the viable region in its state space^8^, i.e. the set of all states beginning in which the walker can step forever by applying at least one sequence of its push-off inputs for every starting state. The viable regions of walking models with definite swing leg dynamics, including the compass walker studied here, have not yet been explicitly estimated. Conversely, in the nonviable set of states, the walker cannot avoid falls, let alone regulate to achieve task goals, even with the best-possible active push-off control. Therefore, the viable region is also the set within which different motor regulation strategies can be meaningfully compared for their effect on the walker’s ability to avoid falls. Taking step-to-step speed regulation as a model task-level motor regulation strategy^9,16,17^, we estimate the speed-regulated walker’s basins for several target speeds vis-à-vis the viable region. Not only do the speed-regulated walker’s basins occupy large regular regions, but we find that only a small collection of these basins covers nearly the entire viable region itself. Motivated by these results, we propose a hierarchical task switching controller that, at least in principle, allows the walker to avoid falls by appropriately switching between different task-level regulators at each walking step. Our work suggests a possible mechanism by which humans could avoid falls, by exploiting redundancy in previously learned regulation strategies to achieve task goals in any given walking context, including that of responding to a large, unexpected disturbance.

**Figure 1.**
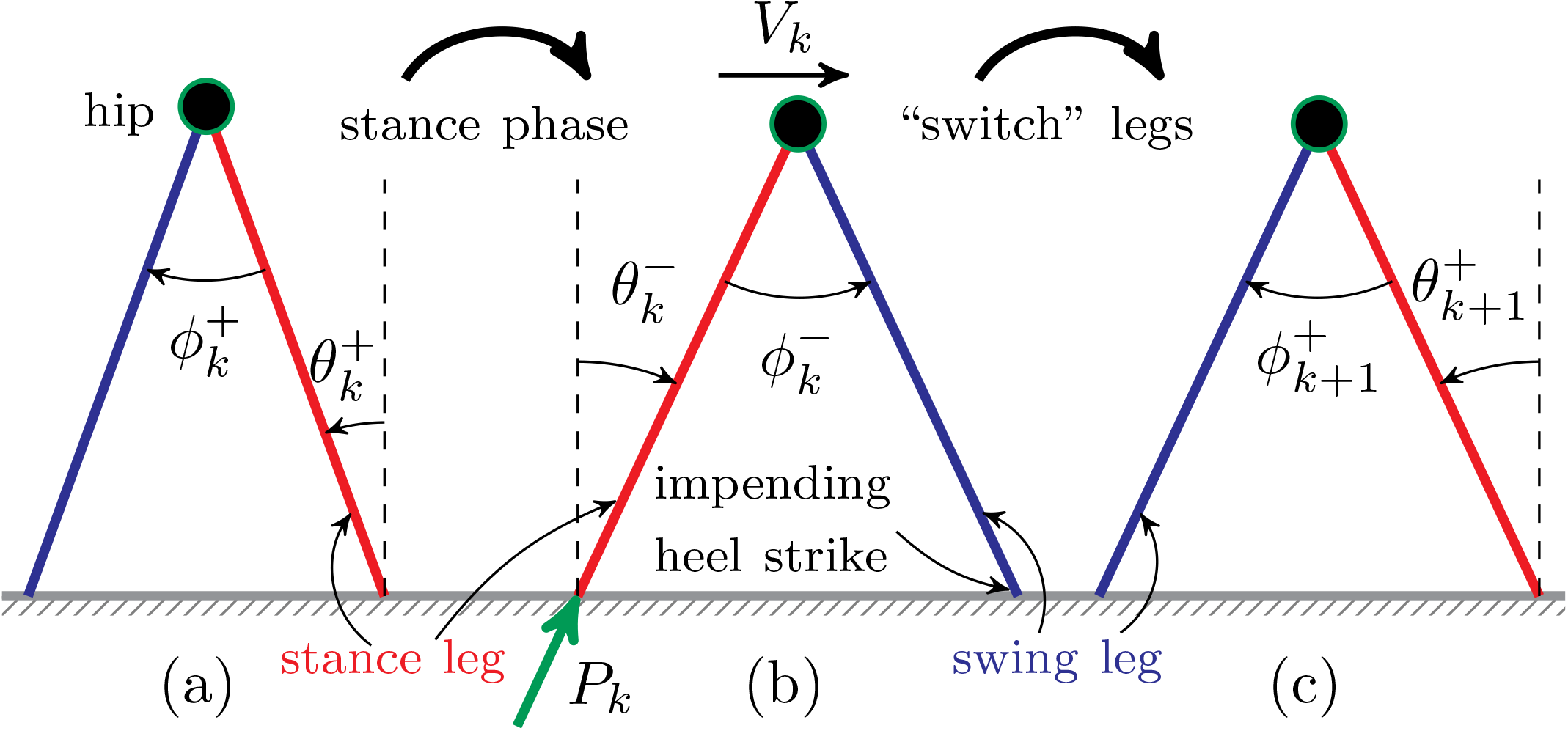
Three snapshots of a 2D powered compass walker^19^ walking on a level ground (step speed, *V*_*k*_): (a) just after *k*^th^, (b) just before (*k* + 1)^st^ and (c) just after (*k* + 1)^st^ heel strike. The walker has straight, massless, stance (*red*) and swing (*blue*) legs, and a mass at the hip (*circle*). The masses at the feet (not shown) are infinitesimally small compared to the hip mass. The push-off impulse, *P*, is applied instantaneously just before heel strike. At the beginning of the *k*^th^ step, the walker’s state in the inertial frame is 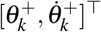.

## Results

We employ the *simplest dynamic walker* that walks on a level surface by means of impulsive push-off actuation, modeling ankle plantar flexion during toe-off in humans^19^ (see “Methods”). The walker’s state, just after heel strike, is fully described by the stance leg angle *θ*^+^ and its angular rate 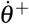, in the inertial frame attached to the stance foot (Fig. 1). We study the walker’s step-to-step dynamics as a *hybrid* Poincaré map, ***F***, over the two-dimensional state space 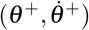 with push-off input *P* (equation 2).

We further impose *viability constraints* on the walker that yield restrictions on its states and inputs (see “Methods”): specifically, the stance foot must remain on the ground throughout the stance phase; the impulsive actuation must be small enough to not lift the walker off the ground when the swing foot’s heel strike is impending, and must be large enough to lift the stance foot off the ground after push-off.

### Where are the compass walker’s dynamics viable?

#### 1 step viable region

Walking motions can start in the feasible region 𝒱_0_of the state space:

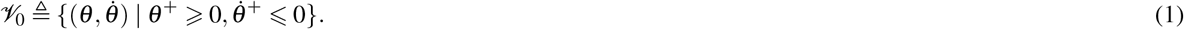

We further restrict *θ*^+^ ⩽ 0.85, which covers the range of stance angles observed in humans.

Previously^17^, we identified the ‘1-step’ region of the powered compass walker (Fig. 1) as the set of states from which the walker can have at least one heel strike. The 1-step region is the wedge-shaped region within 𝒱_0_, demarcated by the curves Ω_low_and Ω_high_(Fig. 2*a*). However, our previous work did not seek to identify the walker’s viability within this region.

**Figure 2.**
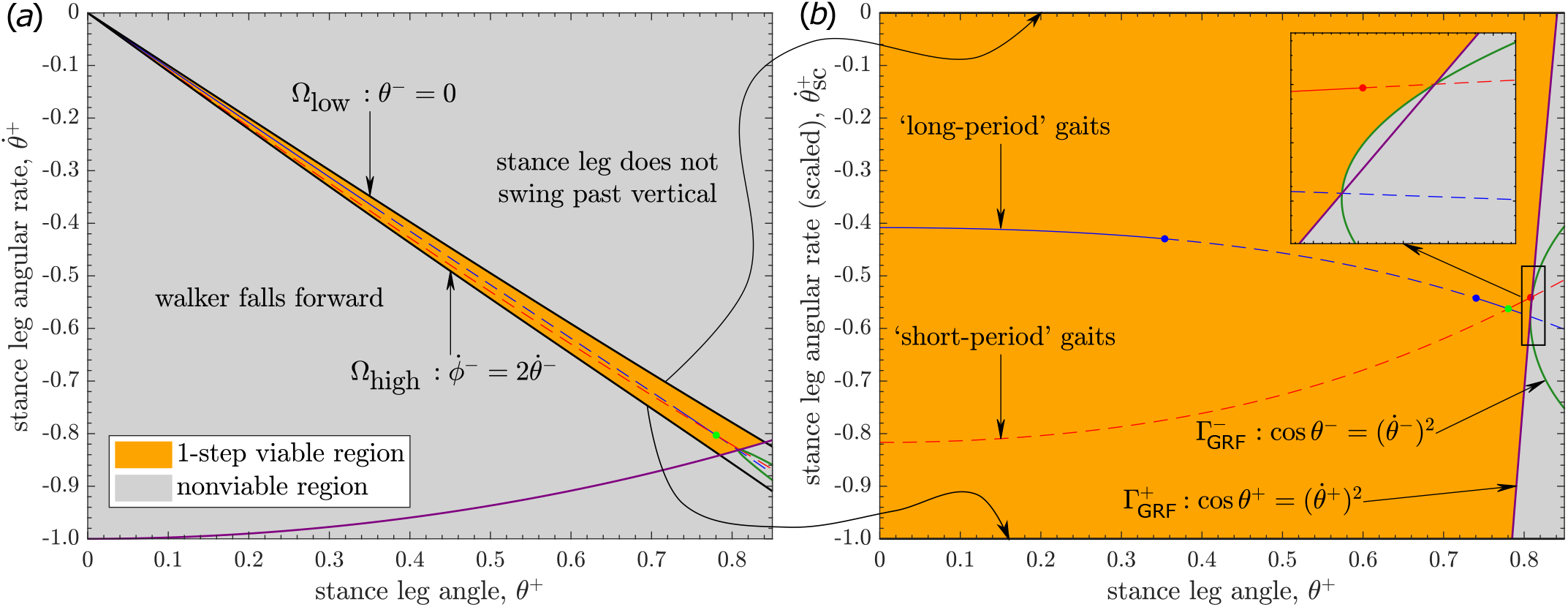
Powered compass walker’s *1-step viable region* 𝒱_1_, i.e. the set of states beginning in which the walker takes at least one step while remaining viable, bounded by the curves Ω_low_, Ω_high_, 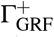, and 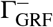: (*a*) In the wedge-shaped region in the middle (‘1-step’ region^17^), demarcated by the curves Ω_low_and Ω_high_, the walker has at least one heel strike though it may not necessarily maintain a nonnegative GRF at the stance foot. Indeed, in the *nonviable region*, the stance leg either moves too slowly to swing past the vertical, moves too fast so that the walker falls forward, or fails to maintain ground contact throughout the stance phase. Thus, 𝒱_1_is a strict subset of the ‘1-step’ region^17^. (*b*) To better visualize 𝒱_1_, we plot the state space with 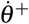 scaled to 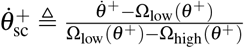 for any given *θ*^+^ ∈ (0, 0.85], so that the new variable 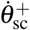 is 0 on the upper Ω_low_curve and takes a value − 1 on the lower Ω_high_curve^17^ (*freehand arrows*). The period-1 gaits of the walker, viz. ‘long-period’ and ‘short-period’ gaits that repeat every step, along with their open-loop stability, are as in our previous work^17^: *solid lines* show open-loop-stable gaits, while *broken lines* depict open-loop-unstable gaits. *Inset* shows zoomed-in area where 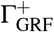, and 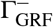 intersect each other and the period-1 gaits.

Here, we find the viable subset of the 1-step region, i.e. the *1-step viable region* 𝒱_1_, by imposing viability constraints on the walker’s dynamics (equation 2). We visualize 𝒱_1_in a scaled state space (Fig. 2*b*), which we introduced previously^17^.

The nonnegativity constraint of the ground reaction force (GRF) at the stance foot yields two curves, 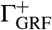 and 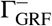 defined by the equalities in equation (5). Each of these curves partition the 1-step region into viable and nonviable sets. Specifically, the walker’s stance foot maintains contact with the ground throughout when initialized from states on the sides of both 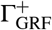 and 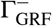 that contain the origin (0, 0) (Fig. 2*b*).

The actuation limits (equation 8) by themselves do not further limit the push-off-powered walker’s viability over a single step. Consequently, the walker’s 1-step viable region 𝔙_1_is bounded by only four curves, viz. Ω_low_, Ω_high_, 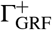 and 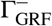 (Fig. 2).

Also shown in Fig. 2 are the walker’s period-1 gaits, i.e. gaits that repeat every step. The walker admits families of ‘long-period’ and ‘short-period’ gaits^6,19^, which are fixed points 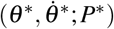 of the map ***F*** (equation 2), for a given *P*^*^. While the long- and short-period gaits admit distinct step times and contrasting open-loop stability as *θ*^+^→ 0^20^, their curves^17^ intersect at (0, 0) in the original state space (Fig. 2*a*). In contrast, in the *scaled* state space (Fig. 2*b*), those very gaits remain bounded away from each other as *θ*^+^ → 0, which facilitates further analysis.

#### Viability kernel: ∞-step viable region

While the walker can definitely take a step in the 1-step viable region 𝒱_1_(Fig. 2), it is not guaranteed to walk forever, even with the best-possible push-off control. This is because the walker’s state after taking a step need not remain in 𝒱_1_, but is only guaranteed to lie in 𝒱_0_(equation 1). We therefore identify the largest closed subset 𝒱 of 𝒱_1_in which the walker can remain viable forever, i.e. for an infinite number of walking steps. That is, for any state 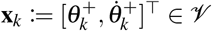, there exists at least one push-off input *P*_*k*_such that 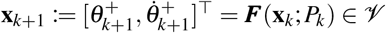, satisfying viability constraints. The set 𝒱 is thus the ∞*-step viable region* or the *viability kernel*^8,15^ of the powered compass walker. It also follows that 𝒱 is the largest *positively invariant* set (i.e. invariant in forward time)^21^ under the walker’s closed-loop dynamics, i.e. with state-dependent push-off input. Alternatively, 𝒱 is the largest *controlled-invariant* set^21^ of the push-off-powered compass walker. Outside 𝒱, the states are *nonviable* as no sequence of push-offs can prevent the walker from eventually failing (i.e. either violating at least one of the viability constraints or falling).

We employed the viability kernel algorithm^8^ that avoids brute-force computation by utilizing the positive invariance property of 𝒱 for its estimation (see “Methods”). Our implementation of that algorithm converged after 18 iterations so that the set 𝒱_18_, i.e. the 18-step viable region where the walker can take at least 18 steps, is the final estimate of the ∞-step region 𝒱 (Fig. 3*a*) to within the resolution of the grid on the state space.

**Figure 3.**
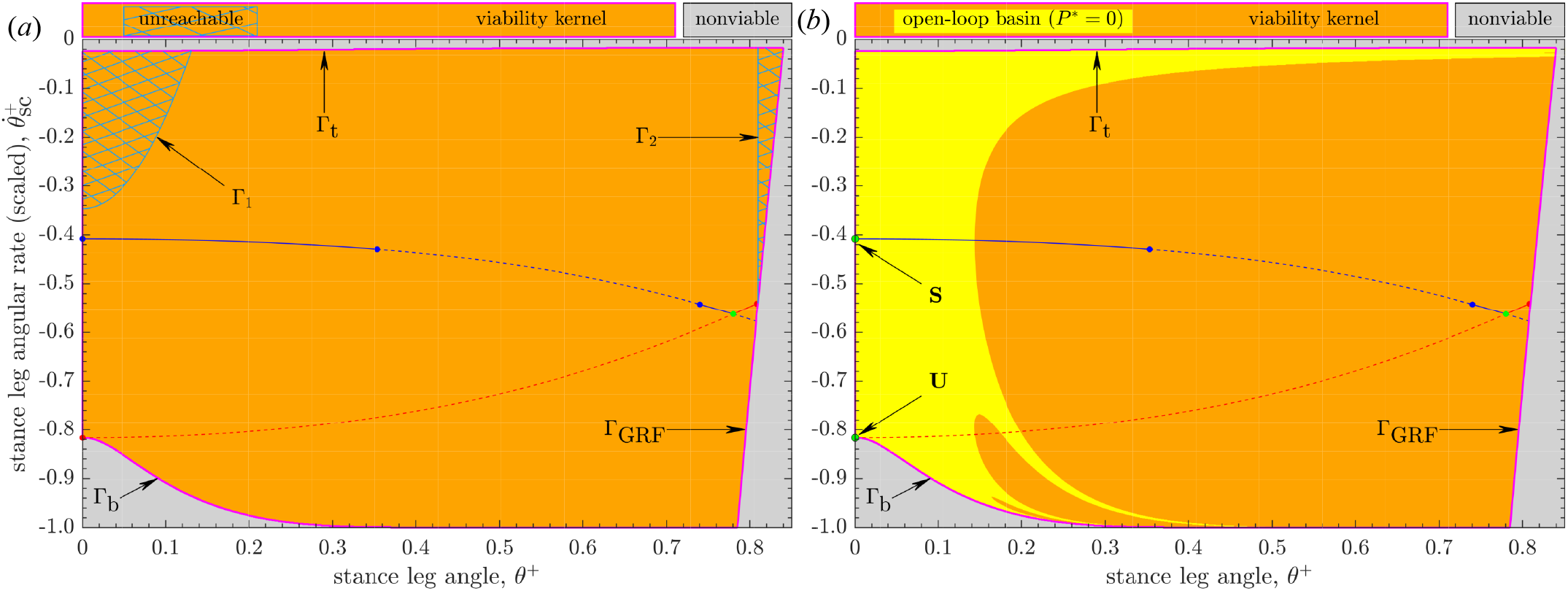
The ∞-step viable region or the *viability kernel* 𝒱 in the scaled state space (Fig. 2*b*) of the powered compass walker, numerically approximated via Algorithm 1 as the 18-step viable region 𝒱_18_after convergence on a grid. The set 𝒱, bounded by the curves {Γ_b_, Γ_t_, Γ_GRF_}, is a strict subset of the 1-step viable region 𝒱_1_(Fig. 2): indeed, states within 𝒱_1_(or ‘1-step’ region^17^) that are either below Γ_b_or above Γ_t_are *nonviable*. The boundary Γ_GRF_is common to both 𝒱 and 𝒱_1_. The curves of long- and short-period gaits are from Fig. 2. (*a*) The two *unreachable* subsets (*hatched* regions) of 𝒱, demarcated by the boundaries Γ_1_and Γ_2_, cannot be traversed by the walker. (*b*) The *open-loop basin* of attraction with zero push-off (*P*^*^ = 0) is a subset of 𝒱, within which trajectories approach the steady-state long-period gait **S** at 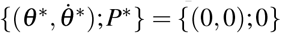. The basin boundaries form the stable set^17,36^ of the unstable short-period gait **U** (a saddle point) at 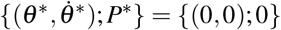 and also contain the boundaries 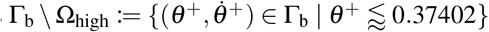 and Γ_t_of 𝒱.

We estimated the areas of different regions in the original state space using the composite Simpson’s rule. The ∞-step viable region 𝒱 (Fig. 3*a*) occupies ≈ 97.46% area of the 1-step viable region 𝒱_1_(Fig. 2): This indicates that the walker’s push-off can be chosen to make it walk forever beginning in almost all states for which it can have a legitimate heel strike (equation 4).

#### Unreachability within the viability kernel

We found the *unreachable* subset 𝒱_UR_of 𝒱 that cannot be traversed by the walker’s trajectories (see “Methods”). The set 𝒱_UR_consists of two disjoint subsets of 𝒱, together occupying ≈ 2.47% of its area (Fig. 3*a*). As expected, the walker’s period-1 gaits lie entirely within the reachable subset of 𝒱.

Evidently, any walking task or target that would require the walker to traverse such unreachable sets cannot be achieved. The walker’s state can end up in 𝒱_UR_due to external disturbances or can be initialized within it; however, its state immediately (i.e. in one step) leaves 𝒱_UR_under the walker’s dynamics (equation 14).

#### Viability kernel boundaries

The viability kernel algorithm guarantees that the trajectories of nonviable grid-point states (Fig. 3) cannot enter the viability kernel 𝒱 while those originating in the interior of 𝒱 always remain in it. However, 𝒱 is a closed set^8^, so states on its boundary must also satisfy the positive invariance property (equation 13): That is, the boundary of 𝒱 can be mapped into itself or into the interior of 𝒱 ^22^, provided appropriate input push-offs are chosen.

The boundary of 𝒱 is a union of three curves: Γ_b_, Γ_t_and Γ_GRF_(Fig. 3). Our numerical results indeed show the positive invariance of the estimated boundaries of 𝒱, which leads to their validation via the mathematical theory of dynamical systems (see “Methods”).

### Task-level regulation, global stability, and fall avoidance

No strategy can avoid falls for states outside the viability kernel 𝒱. Conversely, the walker can walk forever inside 𝒱 by employing any one of infinitely many appropriate sequences of push-offs. However, the region 𝒱 itself exists and is independent of any given control strategy the walker might choose. Thus, it is meaningful to assess the performance of different *specific* control strategies in avoiding falls only for the trajectories starting within 𝒱.

We are particularly interested in motor regulation templates, i.e. empirically motivated models of how humans manipulate task-level observables on a step-to-step basis^10,16^. As a model task-level regulation strategy, we here specify experimentally informed step-to-step speed regulation^9,16^ on the walker (Fig. 1): see “Methods”. Specifically, we pick a push-off impulse at each step by minimizing the squared discrepancy between the speed *V* at the next step and its desired target value *V* ^*^, chosen *a priori*.

#### Global stability under task-level regulation: basins of attraction

Previously^17^, we demonstrated a functional connection between task-level motor regulation and the walker’s ability to reject large disturbances, i.e. its global stability. The maximal attainable global stability for the walker, capable of applying arbitrary sequence of push-offs within its actuation limits, is, indeed, its viability. Therefore, we assess a walker’s global stability via the basins of attraction of its steady-state gaits in the state space vis-à-vis the viability kernel 𝒱 (Fig. 3).

As in our recent work^17^, we numerically estimated basins by simulating the open-loop (*P*_*k*_:= *P*^*^) walker’s trajectories for 50 steps and those of the speed-regulated (equation 16) walker for 25 steps, starting from every state on the same grid that we used for the estimation of 𝒱. The walker’s trajectories that fail to satisfy viability constraints are not considered part of its basins.

The open-loop basins (Fig. 4) are significantly smaller in area than those of the speed-regulated basins (Fig. 5). Moreover, the geometric structure of the open-loop basins becomes more intricate as push-off impulse *P*^*^ increases, with a growing number of disjoint boundaries (see^17^ for a discussion of the aspect of the noninvertibility structure of the open-loop basins). In contrast, the speed-regulated walker’s basins occupy large areas within 𝒱 and are highly regular (Fig. 5): their boundaries are given by level curves of the form *θ* ^−^ = constant^17^ and/or shared with the boundaries of 𝒱 themselves.

**Figure 4.**
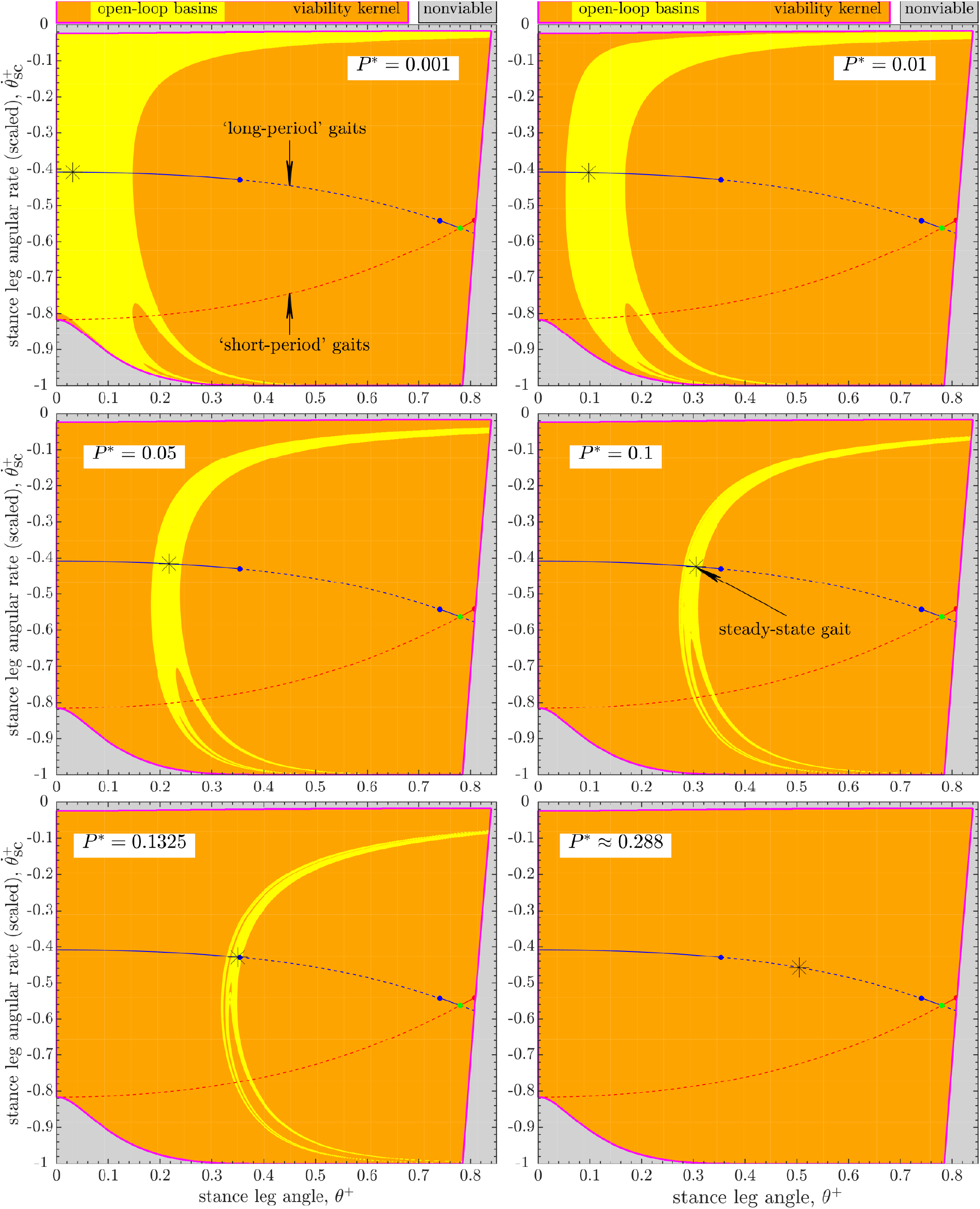
Evolution of the open-loop (*P*_*k*_:= *P*^*^, constant) walker’s basin of attraction within the viability kernel 𝒱 of its long-period steady-state gait (*) with increasing push-off impulse *P*^*^. Compare to Fig. 5. The open-loop basin at *P*^*^ = 0.001 is similar in structure to that with no push-off (*P*^*^ = 0) that shares boundaries with 𝒱 (Fig. 3*b*). As *P*^*^ increases, the basins shift to the right within 𝒱, while shrinking progressively for *P*^*^ ⩾ 0.01. The first period-doubling bifurcation occurs at *P*^*^ ≈ 0.13571, so that the open-loop basin at *P*^*^ ≈ 0.288 is empty. Basin areas (% of the area of 𝒱) for increasing *P*^*^: {8.32, 8.36, 5.38, 3.11, 1.46, 0}%.

**Figure 5.**
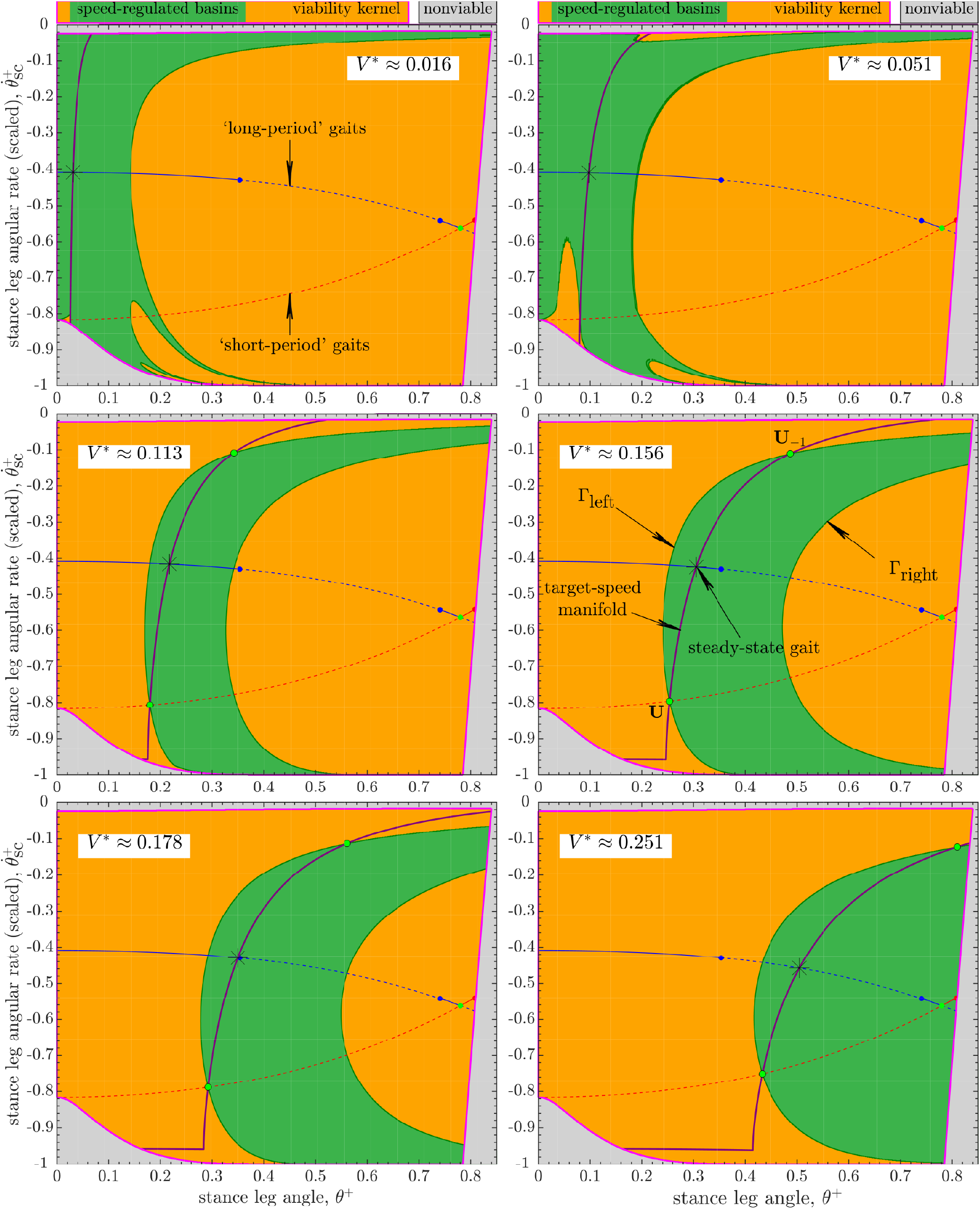
Evolution of the speed-regulated walker’s basin of attraction within the viability kernel 𝒱 of its long-period steady-state gait (*) with increasing target step speed *V* ^*^ (or, push-off impulse *P*^*^∈{ 0.001, 0.01, 0.05, 0.1, 0.1325,≈ 0.288). Compare to Fig. 4. For *V* ^*^ ⩾ 0.113, the basins are highly regular regions delimited by the level curves (Γ_left_and/or Γ_right_of the form {*θ* ^−^ = constant}) and the boundaries of 𝒱 themselves. The geometric structure and size of the speed-regulated basins at *V* ^*^ ∈ {≈ 0.016, ≈ 0.051} is affected by viability constraints, specifically, actuation limits (equation 8): the basin at *V* ^*^ ≈ 0.016 is similar in structure to the open-loop basin for *P*^*^ ∈ {0, 0.001} (Figs. 3 and 4). Basin areas (% of the area of 𝒱) for increasing *V* ^*^: {8.12, 11.62, 20.59, 36.62, 44.83, 54.13}%.

In Fig. 6, we compare the normalized areas of the basins of attraction within the viability kernel 𝒱 for the open-loop and speed-regulated walkers for target speeds *V* ^*^ ⪅0.38301 (or, push-offs *P*^*^ ⪅0.79478), leading up to the transcritical bifurcation^17^.

**Figure 6.**
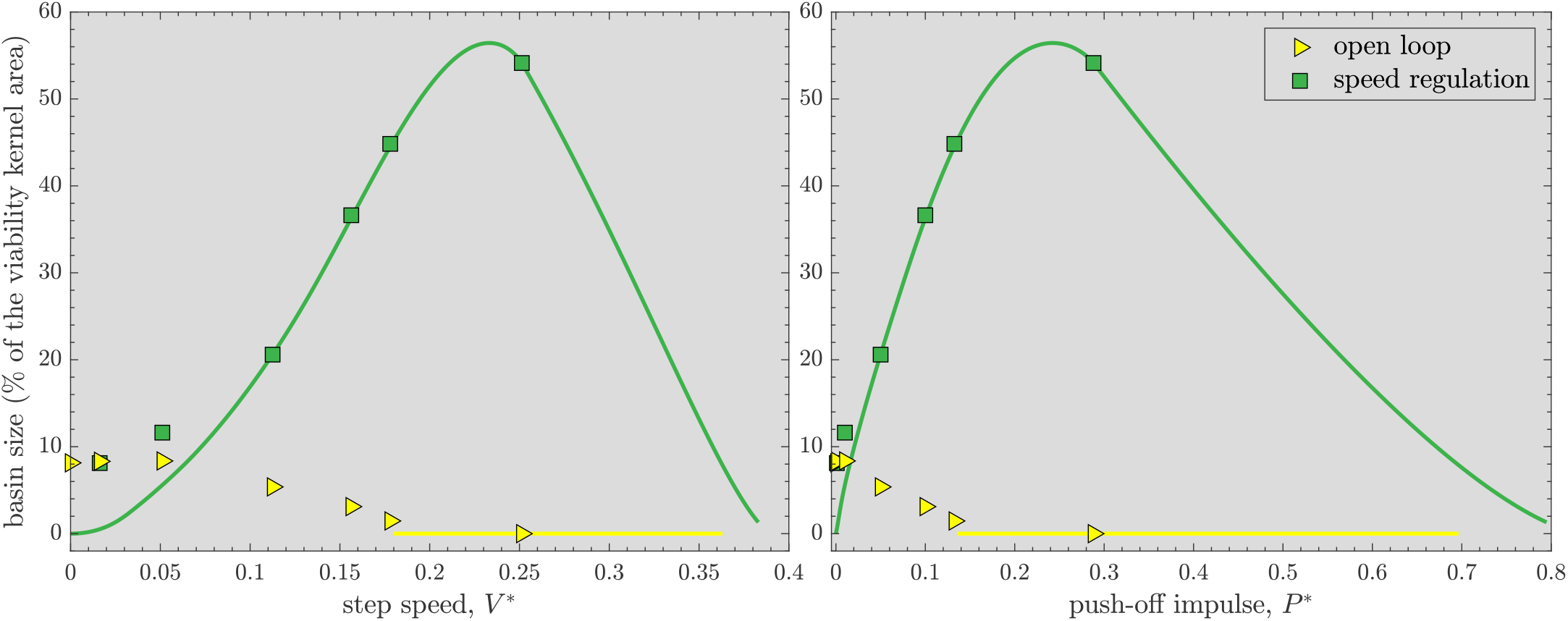
Evolution of the basin sizes [percent of the area of the viability kernel 𝒱 (Fig. 3)] for the open-loop and speed-regulated walkers. Both *V* ^*^ and *P*^*^ correspond to the steady-state long-period gait. The *markers* denote sizes of the basins estimated via simulations (Figs. 3, 4, and 5). The *solid line* for *speed regulation* denotes analytical approximations of basin sizes when basin boundaries can be predicted (either as level curves and/or coinciding with the boundaries of 𝒱): these predictions match simulations well except when actuation limits significantly affect the basin structure at low speeds or push-offs. The *solid line* for *open loop* corresponds to unstable gaits so that their basins are empty.

The open-loop walker’s basin shrinks significantly as *P*^*^ increases from 0.01 to 0.1325, before the long-period gait loses open-loop stability at *P*^*^≈ 0.13571 via a period-doubling bifurcation. The open-loop basin occupies a maximum of ≈ 8.4% area of 𝒱 at *P*^*^ = 0.01. In comparison, the speed-regulated walker’s basin of its long-period gait grows with speed until it achieves its maximum size, ≈ 56.4% area of 𝒱, at *V* ^*^ ≈ 0.23308 (*P*^*^ ≈ 0.24214) before shrinking significantly at higher speeds.

#### Viability via hierarchical task switching control

The open-loop basins in Fig. 4 together occupy only ≈ 20.36% of the area of the viability kernel 𝒱 with many hard-to-fill gaps in between. Furthermore, we estimate that all of the open-loop basins, corresponding to *P*^*^ values of all the long-period gaits, together can cover no more than 40% of the area of 𝒱.

Conversely, the task-level speed regulator, while achieving the specified goal of maintaining a target speed at each step, allows the push-off powered compass walker to reject a large range of external disturbances, despite not being designed to do so^17^. The speed-regulated walker’s basins occupy large, regular regions of 𝒱 for a range of target speeds *V* ^*^ (Fig. 6). Furthermore, as we show in Fig. 7, only five of the speed-regulated walker’s basins from Fig. 5 almost fully cover 𝒱. Thus, starting from almost every state in 𝒱, as might occur from an external disturbance, there is at least one task-level speed regulator (or *V* ^*^) that allows the walker to avoid falls as long as the state trajectory remains within the corresponding basin. Additionally, since a set of target speeds *V* ^*^ can be chosen so that any two adjacent speed-regulated basins overlap (as in Fig. 7), there is flexibility to switch between the corresponding regulators immediately 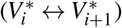 provided the walker’s state lies within the basin intersection. Thus, this suggests that task-level speed regulation, unlike open-loop dynamics, could, at least in principle, be used to keep the walker viable for almost all states in 𝒱, i.e. allowing it to avoid falls forever, in response to any disturbance that does not push the system entirely out of 𝒱. For example, a plausible task switching controller could appropriately switch target speed at each step to one of the five values 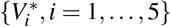, as in Fig. 7, so that the walker can move from one speed-regulated basin to another without falling. We posit that a similar adaptive hierarchical control/regulation strategy exists in human walking and provides a key mechanism used to avoid falling.

**Figure 7.**
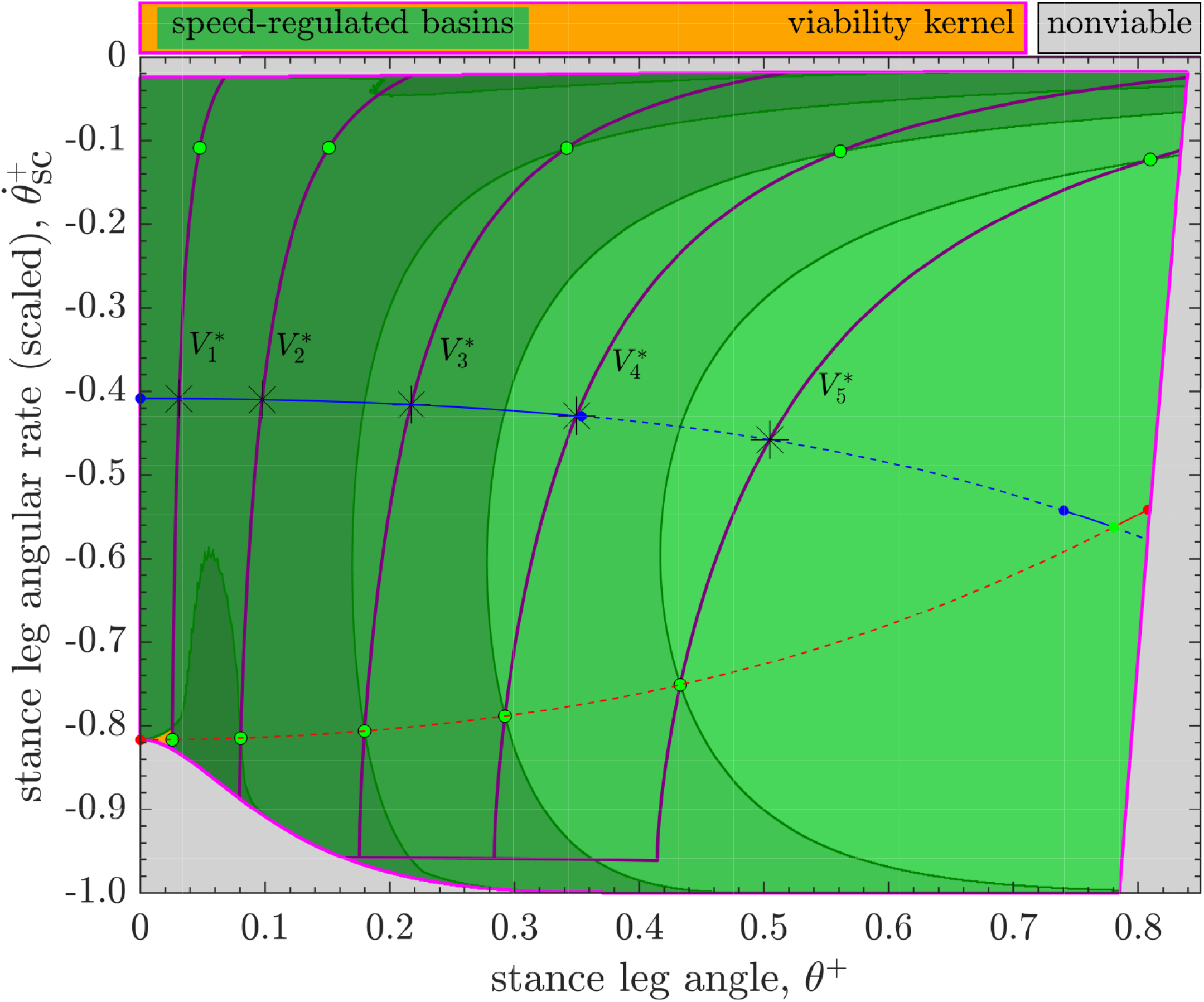
Five of the speed-regulated walker’s basins from Fig. 5 of its long-period steady-state gait (*) corresponding to the target step speeds,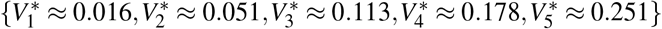. These five basins together cover *>* 99.99% of the area of the viability kernel. Also, any two adjacent basins have some overlap with each other.

To further elucidate the function of such task switching control, we consider a scenario where the walker experiences a large disturbance while maintaining some desired speed 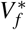. Let the state of the walker immediately after the disturbance lie within the viability kernel: **x**_*k*_∈ 𝒱 at the *k*^th^ walking step. Moreover, we assume that 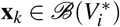, where 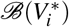 is the task-regulated basin corresponding to some suitably chosen intermediate target speed 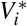. The walker then applies a push-off 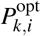 to achieve the target value 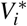 at the next step (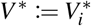 in equation 15). We construct a possibly minimal set, **V**_*p*_, of all such target speeds 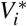 such that the corresponding set of speed-regulated basins together can cover the viability kernel: 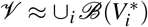 (Fig. 7). Thus, specifying such a hierarchical control strategy amounts to specifying a suitable sequence of ‘task switches’, i.e. target speeds 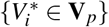 at each step, for the regulators. Such task switching control can, in principle, allow the walker to get back to its original task goal 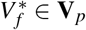 while remaining viable throughout its post-disturbance recovery phase: at the (*k* + 1)^st^ walking step, the walker’s state **x**_*k*+1_not only belongs to 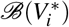 but also to 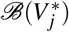 for some 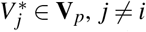, by design. Once the walker’s state trajectory enters the basin 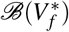 corresponding to the original task, the relevant speed regulator to achieve that task goal is switched back on for subsequent walking steps, until the next large disturbance is encountered. Thus, following such a strategy of switching between a small set of target speeds, the walker could walk forever while overcoming a wide range of large disturbances. Furthermore, because task switching is not mechanical, it is not affected by the the walker’s inertial properties. Thus, the time scale of task switching in humans would be limited not by mechanics proper, but by the speed of processes in the nervous system related to perception, motor activation, and cognition (particularly executive function). Therefore, task switching can, in principle, be accomplished almost instantaneously. This has obvious benefits for recovering from sudden, unexpected disturbances.

## Discussion

We studied the simplest dynamic walker’s viability, i.e. its ability to avoid falls forever by applying an appropriate sequence of push-off inputs. Specifically, for the push-off-powered compass walker^19^ we estimated the viability kernel 𝒱 in its state space^8^ and verified that our numerical results are consistent with the mathematical theory of dynamical systems. We found that the walker’s push-off can be chosen to avoid falls forever beginning in almost all states that allow the walker to have a heel strike. Moreover, greater than 97% of the states within 𝒱 remain reachable via push-off inputs, indicating a high degree of maneuverability of the viable walker.

We posited that humans could remain viable, i.e. avoid falls forever, while carrying out specific goal-directed walking tasks via a hierarchical schema consisting of both control and functionally distinct task-level step-to-step regulation of gait observables. As a model task-level motor regulation strategy for the walker, we imposed speed regulation^9,16^ that, as we demonstrated in^17^, greatly enhances the walker’s global stability (large disturbance rejection) compared to open-loop (unregulated) dynamics. Here, however, we assessed the walker’s global stability relative to its viability—its maximal attainable global stability—via the basins of attraction of its steady-state gaits in the state space vis-à-vis 𝒱. This facilitated a direct comparison between open-loop dynamics, task-level regulation, and theoretically best-possible control strategies from the perspective of fall avoidance alone.

We found that the speed-regulated walker’s basins, unlike the open-loop basins (Fig. 4), occupy large, regular regions within 𝒱 (Figs. 5 and 6). Moreover, for a range of target speeds, their boundaries are given by simple level curves and/or are shared with the boundaries of 𝒱 themselves. Furthermore, the speed-regulated basins corresponding to only a few target speeds together nearly cover the entirety of 𝒱 even as any adjacent pairs of such basins overlap in the state space (Fig. 7). Our results thus strongly suggest a potential role of task-level regulation within high-level control strategies that are explicitly geared toward avoiding falls or attaining viability. In light of this, we proposed a high-level, adaptive task switching control strategy that, in principle, maintains viable walking by switching between a small collection of task-level speed regulators corresponding to a few preselected target speeds—’task switches’—at each walking step. However, it is clear that, at least in principle, such task switching controllers could also employ qualitatively different regulators, based on gait observables other than walking speed, or even a combination of such regulators. Thus, our proposed task switching schema is more general than switching via speed regulation alone.

The theoretically best-possible control strategies that guarantee the walker’s viability would perhaps require specifying an entire sequence of push-offs for each different walking trajectory. In comparison, a hierarchical task switching controller seems advantageous from an information transmission and processing perspective: it needs specifying only a sequence of discrete task switches, each belonging to a small predetermined set (or ‘alphabet’). While we are agnostic as to how such hierarchical task switching control could be realized biologically, our results nevertheless suggest that its ‘information cost’ could be relatively low for the nervous system. This is because the cognitive demands of discretely switching between a few (and likely already learned or ‘crystallized’^23,24^) tasks could be substantially lower than estimating/specifying appropriate control inputs ‘from scratch’ at each walking step. This suggests that humans might prefer cognitively less-demanding hierarchical control strategies based on task switching. Indeed, task switching (or ‘set shifting’) is a well-established sub-component of executive function^25^. For older adults, executive function is crucial to their ability to avoid falling and impaired executive function predicts their fall risk^26–28^. Task switching in particular declines in older adults^29^ and predicts both poor balance^30^ and fall history^31^. Our results thus provide direct theoretical support to the idea that the impaired ability to task-switch appropriately and/or quickly enough likely contributes to increased fall risk in older adults.

While our perspective is focussed on goal-directed behavior of biological movement, our results have implications to robotics as well. Indeed, some high-level strategies based on switching between different controllers^32^ or between limit cycles with speed changes have demonstrated improved stability and versatility of bipedal walking^33^ and running^34^ robot models. For multi-degree-of-freedom robot models, it is computationally difficult to map out their viability kernel in the high-dimensional state space. However, the concept of task switching within a hierarchical control/regulation strategy could be potentially employed to enhance the robustness of walking robots and help reduce (or perhaps minimize) their falls.

## Methods

### Simplest dynamic walker

We employ a 2D compass walker (Fig. 1) that walks on a level surface by means of impulsive push-off actuation *P*. The continuous stance phase of this walker is fully unactuated with no foot placement control. This makes it the simplest actuated model having definite swing leg dynamics, unlike 2D inverted pendulum models^15^.

Every forward walking step (Fig. 1) consists of a continuous-time single-support stance phase followed by an instantaneous impulsive double-support phase. Thus, the walker’s step-to-step dynamics are inherently *hybrid*. The walker’s state, just after heel strike, is fully described by the stance leg angle *θ*^+^ and its angular rate 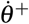, in the inertial frame attached to the stance foot. The walker’s step-to-step dynamics can be studied as a hybrid Poincaré map, ***F*** ≜ [*F*_1_, *F*_2_]^T^, over the two-dimensional state space 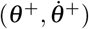 with push-off input *P*_*k*_applied just before heel strike at the end of step *k*^17,19^:

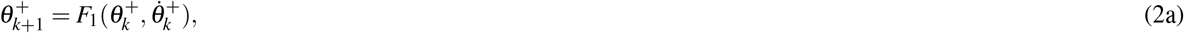

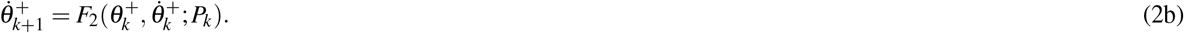

The map ***F*** is *non-invertible*^17^, i.e. any given state of the walker could have zero, one, or more than one preimage under ***F***, even when *P* is fixed. Also, across heel strikes^19^ (Fig. 1):

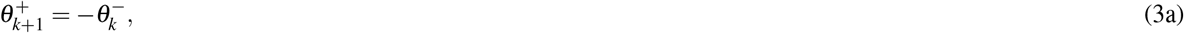

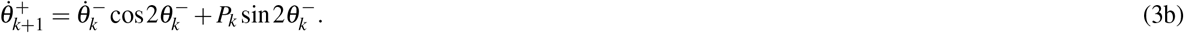

The walker’s heel strike is legitimate (Fig. 1) when:

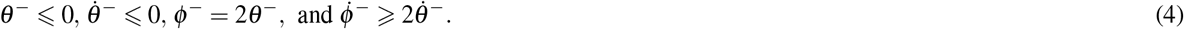

We also assume no slipping at the foot-ground contact.

### Viability constraints

For the walker to remain viable, its stance foot must remain on the ground so that the ground reaction force (GRF) at the stance foot is nonnegative throughout the stance phase. Since this GRF can be smallest either just after or before heel strike, we get two inequality constraints over the state space (Fig. 1):

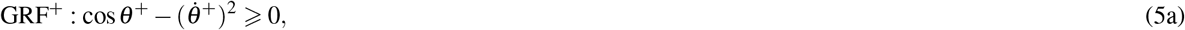

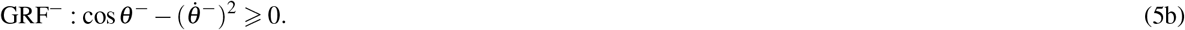

Moreover, this places a state-dependent limit on the maximum push-off, *P*_*k*,max_, since the walker cannot lift off the ground when the swing foot’s heel strike is impending^15^:

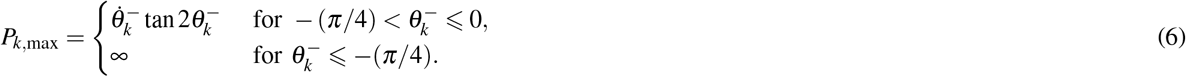

Furthermore, the impulsive actuation cannot apply a braking force, i.e., *P*_*k*_0. Additionally, we assume that the stance foot must lift off the ground after push-off. This places a state-dependent limit on the minimum push-off, *P*_*k*,min_, so that the walking motion can continue (Eqs. 3b and 1):

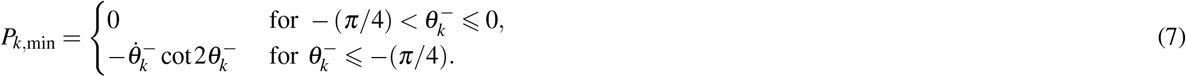

Therefore, the push-off impulse at each step *k* needs to satisfy actuation limits (Eqs. 6 and 7) for the walker to remain viable:

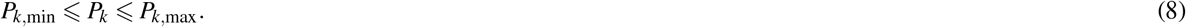

### Numerical estimation of the viability kernel

Estimating the viability kernel 𝒱 for a given actuated dynamical system is in general a non-trivial task, even in low-dimensional state spaces^21^. For instance, 𝒱 is more difficult to estimate than a basin of attraction, another positively invariant set. The trajectories originating in 𝒱 can only be guaranteed to remain in an *as yet unknown* 𝒱 ad infinitum by choosing an appropriate input sequence. Brute-force estimation of 𝒱 requires the computation of many sufficiently long trajectories starting from each state, each of which differ due to distinct control input sequences. If at least one such trajectory satisfies viability constraints, then the corresponding starting state would likely belong to 𝒱. Conversely, the trajectories of basin states approach an attractor that is often *known a priori*. Thus, brute-force estimation of a basin requires only a *single* sufficiently long trajectory starting from each state and a decision as to whether or not it will eventually reach the attractor. A recent study^15^ examined the viability of 2D inverted pendulum models of walking, which have a 1-dimensional state space and a 2-dimensional control input space. In contrast, the system considered here has a 2-dimensional state space and a 1-dimensional control input space.

The viability kernel algorithm [8, pp. 153-154] avoids brute-force computation by utilizing the positive invariance property of 𝒱, which for the walker (equation 2) can be written as:

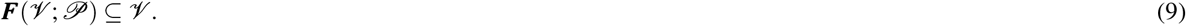

Here, the set ***F***(𝒱; 𝒫) ≜{***F***(**x**; *P*(**x**)) | **x** ∈ 𝒱, *P*(**x**) ∈ 𝒫}, where *P*(**x**) is any appropriately chosen push-off *P* depending on the state 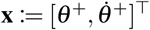, and the set 𝒫 is the collection of all such push-offs matched to states such that the relation (9) holds. The positive invariance property pertains to sets in the state space alone. Since 𝒱 is not known *a priori*, we consider all push-offs within the actuation limits (equation 8) to eliminate 𝒫 from the relation (9).

The dynamics of a powered compass walker, capable of applying any push-off within the actuation limits at each step, is described by a difference inclusion^8^, i.e. a set-valued map 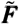 satisfying

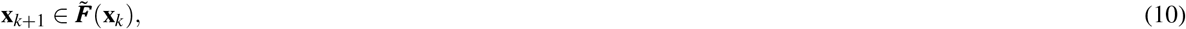

where the set 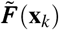 is obtained from equation (2):

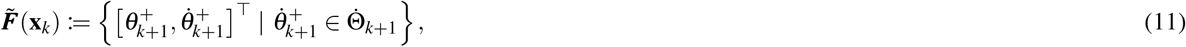

where 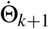 is an interval defined, using equation (3b) and the allowable range of push-offs *P*_*k*_(Eqs. 6 and 7), as:

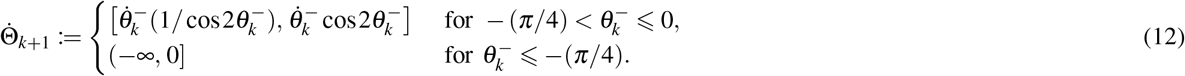

Thus, at step *k*, the set 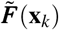 is a vertical line segment, 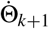, in the state space at 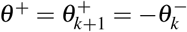 (equation 3a). Therefore, equation (9), expressed solely in terms of states, becomes:

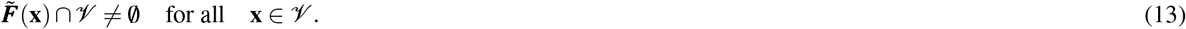

Thus, in principle, one can recursively obtain 𝒱 via the following algorithm:

#### Algorithm 1: discrete viability kernel^8^

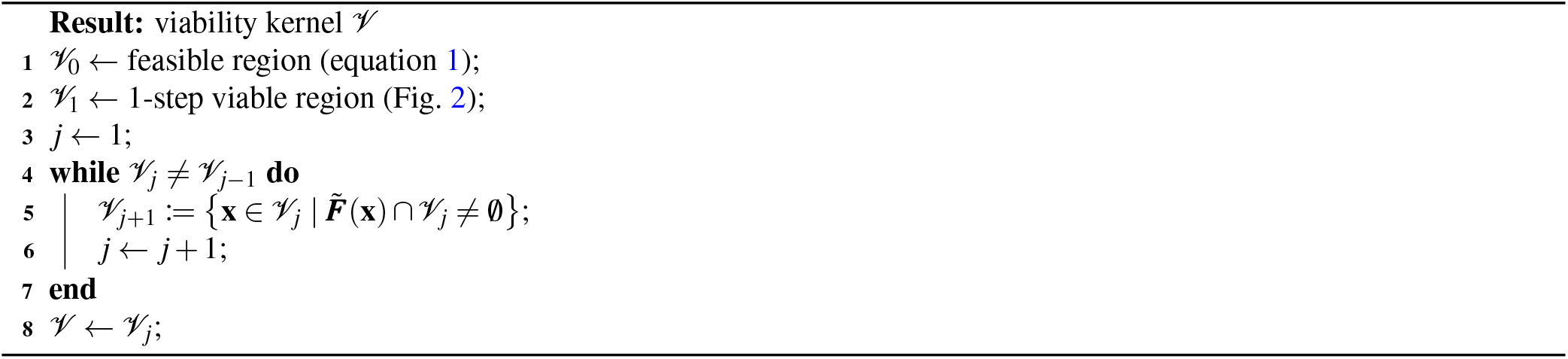

The intermediate estimates {𝒱_*j*+1_⊆ 𝒱_*j*_; *j* = 1, 2, 3, ….} form a nested sequence of *j*-step viable regions, whose limit is the viability kernel: 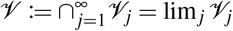. We numerically approximate 𝒱 via a uniform 42500×1002 grid of points over the scaled state space (Fig. 2*b*): Δ*θ*^+^ = 2 ×10^−5^ and 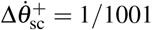. However, all dynamics calculations are carried out in the *original* state space, for the corresponding grid over the wedge-shaped region (Fig. 2*a*).

Algorithm 1 is practically useful if it converges (stops) in a finite (preferably small) number of iterations. This requires an accurate representation of the boundaries of 𝒱_*j*_(at the *j*^th^ iteration) so that the intersection 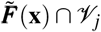 in Algorithm 1 can be found reliably. The boundaries of 𝒱_1_(Ω_low_, Ω_high_, 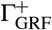 and 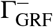 in Fig. 2) are smooth level curves, which we accurately represent via fitted piecewise cubic splines with continuous curvature (matlab’s `spline`). Since sets {𝒱_*j*_; *j* ⩾ 2} are recursively estimated as collections of *j*-step viable grid-point states, their boundaries are not known in closed form. We represent such boundaries by employing shape-preserving piecewise cubic polynomials (matlab’s `pchip`) to reduce potential artifacts (overshoots and oscillations) in the fitted curves over the grid. We passed such fitted boundary curves through nonviable grid-point states tightly enveloping estimates of 𝒱_*j*_so as to avoid accidental removal of viable states during the iterations of Algorithm 1. Our implementation of Algorithm 1 converged at *j* = 18, so that the set 𝒱_18_is the final estimate of the ∞-step region 𝒱 (Fig. 3*a*) to within the grid resolution.

### Unreachable subset of the viability kernel

The image of 𝒱 under 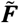, i.e. 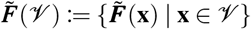, does not cover 𝒱 entirely, so that the *unreachable* subset of 𝒱 is the open set:

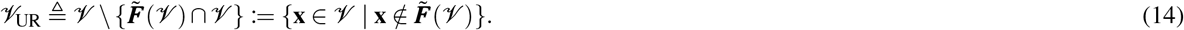

We found that the set 𝒱_UR_consists of two disjoint subsets of 𝒱 demarcated by the boundaries Γ_1_and Γ_2_(Fig. 3*a*). None of the grid-point states in 𝒱 map above the boundary Γ_1_with *P* = 0, and consequently for any *P >* 0 (equation 3b). Studies of non-invertible maps^35,36^ thus suggest that the boundary Γ_1_belongs to a *critical curve* (often denoted as *LC*): the number of preimages of states on opposite sides of *LC* differs by two, which we also found to be the case for Γ_1_.

### Validation of the viability kernel boundaries

The boundary of 𝒱 is a union of three curves: Γ_b_, Γ_t_and Γ_GRF_(Fig. 3). The composite boundary Γ_GRF_itself is a subset of the union of the boundaries 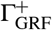 and 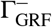 of the 1-step viable region 𝒱_1_(Fig. 2*b*). The boundary Γ_b_smoothly merges with the Ω_high_curve (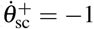 in Fig. 3) at *θ*^+^ ≈0.37402. Thus, Γ_b_is partitioned into two subsets, 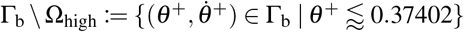 and 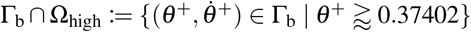. We numerically verified that both Γ_GRF_and Γ_b_⋂Ω_high_can indeed be mapped in the interior of 𝒱.

Furthermore, both boundaries Γ_b_\Ω_high_and Γ_t_map into Γ_b_\Ω_high_after one step of the walker with zero push-off. Moreover, the boundary Γ_b_is tangent to the curve of short-period gaits at the open-loop-unstable gait **U** (a saddle point) at 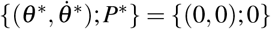 (Fig. 3). These numerical results suggest that Γ_b_\Ω_high_and Γ_t_belong to the *stable set* of the saddle **U**^17, 36, 37^ of the walker’s non-invertible map ***F*** with *P* = 0 in equation (2). Consistently, we found that both Γ_b_\Ω_high_and Γ_t_are contained in the open-loop basin boundaries for *P*^*^ = 0 (Fig. 3*b*), which constitute a stable set^17^. Since the stable set of a saddle is positively invariant, this confirms that the set {Γ_b_\ Ω_high_} ∪ Γ_t_is also positively invariant.

### Step-to-step speed regulation as a model task-level regulation strategy

We pick a push-off impulse at each step based on the discrepancy between the speed *V* at the next step that depends on the walker’s current state and its desired target value *V* ^*^, chosen *a priori*. Thus, at step *k*, the smallest push-off, 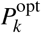, that minimizes the next-step quadratic cost is^17^:

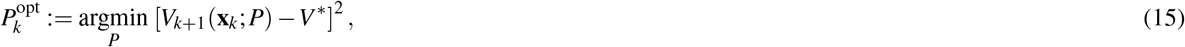

where 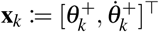 is the walker’s state at the beginning of step *k*. Then, the speed-regulated walker applies push-off *P*_*k*_that satisfies actuation limits (equation 8) at step *k*:

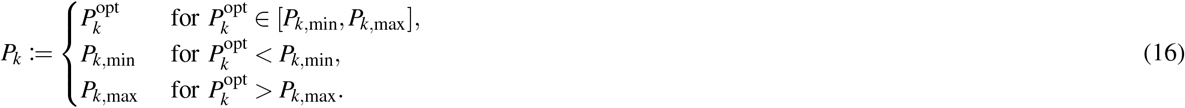

We note that this speed regulation strategy (equation 15) does not explicitly utilize the location of the boundaries of 𝒱 (Fig. 3) to infer 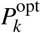.

The push-off 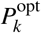 places the speed-regulated walker’s state **x**_*k*+1_on the target-speed manifold—a *goal equivalent manifold*^38^—that is a piecewise-smooth curve in the two-dimensional state space^17^, defined by:

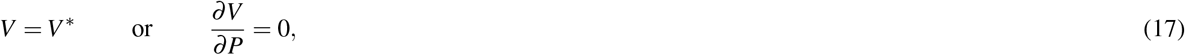

whenever 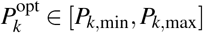 (equation 8). We efficiently simulated the speed-regulated walker’s trajectories by precomputing the target-speed manifold by solving equations (17) using numerical continuation^17^.

## Acknowledgements

This work was supported by the U.S. National Institutes of Health (grant no. R01-AG049735).

## Author contributions statement

All authors participated in the design of the study. N.S.P. performed model analysis, carried out computer simulations, and drafted the manuscript in consultation with J.P.C. The manuscript was reviewed, edited, and prepared for submission by all authors.

## References

1. Adolph, K. E. et al. How do you learn to walk? thousands of steps and dozens of falls per day. Psychol. Sci. 23, 1387–1394, DOI: 10.1177/0956797612446346 (2012).

2. Burns, E. R., Stevens, J. A. & Lee, R. The direct costs of fatal and non-fatal falls among older adults — united states. J. Saf. Res. 58, 99–103, DOI: 10.1016/j.jsr.2016.05.001 (2016).

3. Burns, E. & Kakara, R. Deaths from falls among persons aged ≥ 65 years — united states, 2007–2016. MMWR. Morb. Mortal. Wkly. Rep. 67, 509–514, DOI: 10.15585/mmwr.mm6718a1 (2018).

4. Ambrose, A. F., Paul, G. & Hausdorff, J. M. Risk factors for falls among older adults: A review of the literature. Maturitas 75, 51–61, DOI: 10.1016/j.maturitas.2013.02.009 (2013).

5. Hobbelen, D. G. & Wisse, M. Limit cycle walking. In Hackel, M. (ed.) Humanoid Robots, Human-like Machines, chap. 14, DOI: 10.5772/4808 (IntechOpen, Rijeka, 2007).

6. McGeer, T. Passive Dynamic Walking. Int. J. Rob. Res. 9, 62–82, DOI: 10.1177/027836499000900206 (1990).

7. Todorov, E. & Jordan, M. I. Optimal feedback control as a theory of motor coordination. Nat. Neurosci. 5, 1226–1235, DOI: 10.1038/nn963 (2002).

8. Aubin, J.-P. Viability Theory (Birkhäuser, Boston, MA, 2009).

9. Dingwell, J. B., John, J. & Cusumano, J. P. Do humans optimally exploit redundancy to control step variability in walking? PLoS Comput. Biol. 6, e1000856, DOI: 10.1371/journal.pcbi.1000856 (2010).

10. Dingwell, J. B. & Cusumano, J. P. Humans use multi-objective control to regulate lateral foot placement when walking. PLoS Comput. Biol. 15, 1–28, DOI: 10.1371/journal.pcbi.1006850 (2019).

11. Bernstein, N. A. The co-ordination and regulation of movements (Pergamon Press, Oxford, 1967).

12. Wong, A. L. & Haith, A. M. Motor planning flexibly optimizes performance under uncertainty about task goals. Nat. Commun. 8, 14624, DOI: 10.1038/ncomms14624 (2017).

13. Wieber, P.-B. Constrained dynamics and parametrized control in biped walking. In International Symposium on Mathematical Theory of Networks and Systems (Perpignan, France, 2000).

14. Wieber, P.-B. On the stability of walking systems. In International Workshop on Humanoid and Human Friendly Robotics (Tsukuba, Japan, 2002).

15. Zaytsev, P., Wolfslag, W. & Ruina, A. The boundaries of walking stability: Viability and controllability of simple models. IEEE Trans. Robot. 34, 336–352, DOI: 10.1109/TRO.2017.2782818 (2018).

16. Dingwell, J. B. & Cusumano, J. P. Identifying stride-to-stride control strategies in human treadmill walking. PLOS ONE 10, e0124879, DOI: 10.1371/journal.pone.0124879 (2015).

17. Patil, N. S., Dingwell, J. B. & Cusumano, J. P. Task-level regulation enhances global stability of the simplest dynamic walker. J. R. Soc. Interface 17, 20200278, DOI: 10.1098/rsif.2020.0278 (2020).

18. Full, R. & Koditschek, D. Templates and anchors: neuromechanical hypotheses of legged locomotion on land. J. Exp. Biol. 202, 3325–3332 (1999).

19. Kuo, A. D. Energetics of Actively Powered Locomotion Using the Simplest Walking Model. J. Biomech. Eng. 124, 113–120, DOI: 10.1115/1.1427703 (2001).

20. Garcia, M., Chatterjee, A., Ruina, A. & Coleman, M. The Simplest Walking Model: Stability, Complexity, and Scaling. J. Biomech. Eng. 120, 281–288, DOI: 10.1115/1.2798313 (1998).

21. Blanchini, F. Set invariance in control. Automatica 35, 1747–1767, DOI: 10.1016/S0005-1098(99)00113-2 (1999).

22. Blanchini, F. & Miani, S. Set-Theoretic Methods in Control (Springer International Publishing, Switzerland, 2015), 2nd edn.

23. Tumer, E. C. & Brainard, M. S. Performance variability enables adaptive plasticity of ‘crystallized’ adult birdsong. Nature 450, 1240–1244, DOI: 10.1038/nature06390 (2007).

24. Grafton, S. T. Malleable templates: reshaping our crystallized skills to create new outcomes. Nat. Neurosci. 11, 248–249, DOI: 10.1038/nn0308-248 (2008).

25. Miyake, A. et al. The unity and diversity of executive functions and their contributions to complex “frontal lobe” tasks: A latent variable analysis. Cogn. Psychol. 41, 49–100, DOI: 10.1006/cogp.1999.0734 (2000).

26. Mirelman, A. et al. Executive function and falls in older adults: New findings from a five-year prospective study link fall risk to cognition. PLoS ONE 7, e40297, DOI: 10.1371/journal.pone.0040297 (2012).

27. Montero-Odasso, M., Verghese, J., Beauchet, O. & Hausdorff, J. M. Gait and cognition: A complementary approach to understanding brain function and the risk of falling. J. Am. Geriatr. Soc. 60, 2127–2136, DOI: 10.1111/j.1532-5415.2012.04209.x (2012).

28. van Schooten, K. S. et al. Sensorimotor, cognitive, and affective functions contribute to the prediction of falls in old age and neurologic disorders: An observational study. Arch. Phys. Med. Rehabil. 102, 874–880, DOI: 10.1016/j.apmr.2020.10.134 (2021).

29. Levy-Tzedek, S. Changes in predictive task switching with age and with cognitive load. Front. Aging Neurosci. 9, DOI: 10.3389/fnagi.2017.00375 (2017).

30. Redfern, M. S., Chambers, A. J., Sparto, P. J., Furman, J. M. & Jennings, J. R. Inhibition and decision-processing speed are associated with performance on dynamic posturography in older adults. Exp. Brain Res. 237, 37–45, DOI: 10.1007/s00221-018-5394-0 (2018).

31. McKay, J. L., Lang, K. C., Ting, L. H. & Hackney, M. E. Impaired set shifting is associated with previous falls in individuals with and without parkinson’s disease. Gait & Posture 62, 220–226, DOI: 10.1016/j.gaitpost.2018.02.027 (2018).

32. Saglam, C. O. & Byl, K. Switching policies for metastable walking. In 52nd IEEE Conference on Decision and Control, DOI: 10.1109/cdc.2013.6760009 (2013).

33. Veer, S., Motahar, M. S. & Poulakakis, I. Generation of and switching among limit-cycle bipedal walking gaits. In 2017 IEEE 56th Annual Conference on Decision and Control (CDC), DOI: 10.1109/cdc.2017.8264540 (2017).

34. Bhounsule, P. A., Zamani, A. & Pusey, J. Switching between limit cycles in a model of running using exponentially stabilizing discrete control lyapunov function. In 2018 Annual American Control Conference (ACC), DOI: 10.23919/acc.2018.8431123 (2018).

35. Mira, C., Gardini, L., Barugola, A. & Cathala, J.-C. Chaotic Dynamics in Two-Dimensional Noninvertible Maps (World Scientific, Singapore, 1996).

36. England, J. P., Krauskopf, B. & Osinga, H. M. Bifurcations of stable sets in noninvertible planar maps. Int. J. Bifurc. Chaos 15, 891–904, DOI: 10.1142/s0218127405012466 (2005).

37. Mira, C., Fournier-Prunaret, D., Gardini, L., Kawakami, H. & Cathala, J. Basin Bifurcations of Two-Dimensional Noninvertible Maps: Fractalization of Basins. Int. J. Bifurc. Chaos 04, 343–381, DOI: 10.1142/s0218127494000241 (1994).

38. Cusumano, J. P. & Cesari, P. Body-goal variability mapping in an aiming task. Biol. Cybern. 94, 367–379, DOI: 10.1007/s00422-006-0052-1 (2006).

